# Maternal phylogenetic relationships and genetic variation among rare similar phenotype donkey breeds

**DOI:** 10.1101/2020.04.03.022921

**Authors:** Andrea Mazzatenta, Maurizio Caputo, Francesco De Sanctis, Jordi Mirò Roig, Domenico Robbe, Augusto Carluccio

## Abstract

Maternal inheritance is an indispensable aspect in donkey rare breed population biodiversity management and breeding programs. It is a challenge to characterize breeds genetic inheritance using morphology and historical records, we study mtDNA, to overcome those limitations. The mitochondrial DNA (mtDNA) sequencing is a highly informative system to investigate maternal lineages and breed linkage such as molecular evolution and phylogenetic relationships. Martina Franca, Ragusano, Pantesco and Catalonian donkey mtDNA sequencing analyses were used to study intraspecific genetic diversity and population structure, and to reconstruct phylogenetic relations among these geographically isolated breeds.

A wide lost in variability among all breeds emerged. In this scenario, the primeval haplotypes, higher haplogroups variability and larger number of maternal lineages are preserved in Martina Franca and Ragusano. Accordingly, a putative pivotal role in the phyletic relationship is likely for such breeds.

Given the level of endangerment undergone by these breeds, some actions are necessary to ensure their longtime survival and conservation. Improving the reproduction and management of existing populations, clarifying their historic interactions by studying the genetic status of their populations, extending and improving monitoring maternal lineages represent valid options.

## Introduction

The fact that geographically isolated donkey breeds Martina Franca (MF), typical of continental Puglia region; Ragusano (RG), characteristic of Sicily highland; Pantesco (PT), representative of the small Pantelleria highland (85 Km^2^); and Catalonian (CT) distinctive of the Spanish Catalonian region share similar phenotypes is a matter of genetic inheritance/ecological constrain and biodiversity conservation programs concern. Since the three autochthonous Italian breeds are considered endangered (MF and RG) or critically endangered (PT), the interest in genetic biodiversity safeguarding is growing (FAO report; Rizzi et al, 2011).

Therefore, donkey natural history is a matter of concern. The genus *Equus* is the only remaining of the *Equidae* family, incorporating both extant and fossil species (MacFadden, 2005). The non-caballine forms include: African wild ass, *Equus africanus*; zebras *E. quagga* (formerly *E. burchellii*), *E. grevyi* and *E. zebra* (with two subspecies, *E. z. zebra* of South Africa, and *E. z. hartmannae* of Namibia and Angola); and the Asian wild asses *E. kiang* and *E. hemionus*, with various recognized subspecies *E. h. kulan* and *E. h. onager* (Kruger et al, 2005; Groves & Grubb, 2011). In such picture, domestic donkey (*E. africanus africanus*) is accepted to be a subspecies of African wild ass (Beja-Pereira et al, 2004; Kimura et al, 2011).

The phylogenetic relationship within breed of the domestic form, however, is not well understood. The historical file state about the Roman Empire domination of the Iberian Peninsula, which lasted about 613 years, from 218 BC to 395 (Piganiol, 1927). At that time, in transport and military operations MF donkey was commonly used, consequently its colonization of Spain was highly presumable. Inversely, later in central southern Italy, Spanish domination has occured for about 148 years (1559-1707 AC), and CT could be introduced. However, phenotypic traits and anthropological records are often insufficient to ascertain the breed history, origin and occurrence of genetic exchange (Clutton-Brock, 1992). Instead, the mitochondrial DNA (mtDNA) properties simplify the understanding, through sequencing, of historical, biogeographic and phylogenetic relationship in intra- and inter-species genetics structure (Bowling et al, 2000). The extrachromosomal mitochondrial genome, unlike the nuclear one, is inherited only through the maternal lineage, it is haploid, and its genes do not recombine (Hutchison et al 1974; Brown et al, 1979). The sequence polymorphism identification, by the clonal nature of its heredity, for domestic animals genetic studies is a unique application (Bowling et al, 2000; Rand, 2001). Therefore, the variation of the D-loop region in mtDNA combined with the lack of recombination produces a highly informative tool for matrilineal relationship studies infer intra-species phylogenetic relationships and characterize intra-breed variation (Vigilant et al, 1989; Brown et al, 1979; Wallace, 2007; Mirol et al 2002; Stoneking et al., 1992). Similarly, studies on mtDNA dog breeds, with greater phenotypic and working variability compared to donkey’s uniformity, have revealed genetic information on their domestication, evolution and hereditary diseases (Verginelli et al, 2005; Mazzatenta et al, 2017).

The mtDNA studies of equine breeds were addressed to investigate their origin (Vila et al, 2001; Achilli et al, 2012; Cothran et al, 2005; Kavar & Dovc, 2008; Lippold et al, 2011; Cieslak et al, 2010; Jansen et al, 2002; Guastella et al, 2011); to track breed migration and distribution by comparing maternal lines in different populations (Kivisild et al, 2004; Matisoo-Smith & Robins, 2009). Further, the donkey complete mitochondrial genome sequence was greatly informative (Xu et al, 1996; Luo et al 2011) and was used to date the divergence with horse, ranging between 8 to 10 MYA, which is earlier than both paleontological data (Simpson, 1951; Lindsay et al, 1980) and equids restriction endonuclease analysis proposed (George & Ryder, 1986).

Fascinating, the identification of two lineages in domestic donkey mtDNA termed Clade 1, for Nubian (*Equus africanus africanus*), and Clade 2, for Somali (*Equus africanus somaliensis*), is believed to be the result of two separate domestication events, from two ancestral wild populations existing in the Atbara region and Red Sea Hills (NW Sudan); and in southern Eritrea, Ethiopia and Somalia (Uerpmann, 1991; Vila, 2002; Rossel et al, 2008; Beja-Pereira et al, 2004; Chen et al, 2006). However, the existence of another ancestor of the domestic donkeys, belonging to unrecognized extinct African wild population, has been suggested (Kimura et al, 2011; Kefena et al, 2014).

Genetic studies on the Italian donkey biodiversity are limited, and mainly focused on the variability referred to a protein markers and microsatellites (Ciampolini et al. 2007; Guastella et al. 2007; Matassino et al. 2014; Bordonaro et al. 2012; Colli et al. 2013). Recently, a whole genome sequencing approach (Bertolini et al, 2015) and mtDNA study (Cozzi et al, 2017) were used to study the evolutionary aspects and genetic diversity of Italian donkey populations.

In this scenario, we study the mtDNA D-loop of Italian endangered and critically endangered breeds. mtDNA sequences, polymorphisms (SNPs) and haplotypes were identified and analyzed to investigate the matrilineal assortment within and between asinine breeds with such similar phenotype, as well as the origin and phyletic relationships in other to better asses the management of rare donkey breeds to establishing the proper reproduction and conservation program.

## Materials and Methods

123 salivary samples were collected in eight official breeding stations (Tab. 1), in accordance with the standards for the care and protection of animals used for scientific purposes, Directive 2010/63/EU. The samples collected are from free-range animals with a Certificate of Origin, used to exclude the same maternal descent and select the presumed higher genetic variability. 77 samples (out of 123), divided as follow MF = 27, RG = 22, PT = 8, CT = 19 and an Italian crossbreed, were successfully sequenced (Tab. 1).

Genetic material was collected from saliva using a sterile oral swab, transferred and immortalized on the FTA mini-card and stored in the Multi Barrier Pouches (Whatman Labware Products, UK). The reference material is available at the O.V.U.D. (University Veterinary Hospital) Centre for the breeding of large animals at the Faculty of Veterinary Medicine, University of Teramo Italy. According to the complete donkey mtDNA sequence GenBank X97337 (Xu et al. 1996) two pairs of primers were designed to amplify a 350 bp fragment of displacement loop (D-loop) mtDNA (http://bioinfo.ut.ee/primer3), after extraction from the FTA mini-card, was amplified by the Polymerase Chain Reaction (PCR) performed on 25 μl of reaction volume containing 50 ng of DNA, 2,5 mM of MgCl2, 0.2 mM of each dNTP, 0.5 μM of Per 5’ - CCC AAG GAC TAT CAA GGA AG-3’ and Rew 5’-TTG GAG GGA TTG CTG ATT TC-3’ primer, 1 X PCR Buffer and 1 U of Taq DNA polymerase (Fermentas, Thermo Fischer Scientific). The thermo-amplification cycle was performed using the Mastercycler thermal cycler (Eppendorf, USA) with the following conditions: initial denaturation at 94°C for 5 min followed by 35 cycles of denaturation at 94°C for 30 seconds, heating at 58°C for 30 seconds of extension at 72°C for 30 seconds and final extension at 72°C for 5 minutes.

The raw sequence trace files were checked for the presence of ambiguous bases using software Chromas v. 2.5.1 (http://www.technelysium.com.au/). Sequences were aligned with Muscle. The number of polymorphic sites, parsimony informative and singleton site, number of haplotypes, private and shared haplotypes, haplotype diversity, nucleotide diversity and average number of nucleotide differences were calculated according to Tajima (1983) and Nei (1987) using MEGA7 as well as the maximum parsimony analysis and the maximum composite likelihood method. The median-joining network and principal coordinates analysis (PCoA) were performed with DARwin software (Tamura et al, 2004, Tamura et al, 2012; Kumar et al, 2015; Kumar et al, 2016). The other statistical analysis was performed with Statistica 7.0 StatSoft.

## Results

### Breed analysis

The collection of samples successfully analyzed, from eight certified breeding center (Tab.1), covers the recognized genetic pool range of Martina Franca (MF), Ragusano (RG), Pantesco (PT) and Catalonian (CT) donkey (Tab.1). About 350-bp fragment of the mtDNA displacement loop (D-loop) region was fully sequenced from 77 samples, available through GenBank XXX. A total of 56 haplotypes out of 33 polymorphic sites were found, 14.6% have a frequency higher than 2.7 while 85.4% lower (Fig. 1).

**Fig.1.**
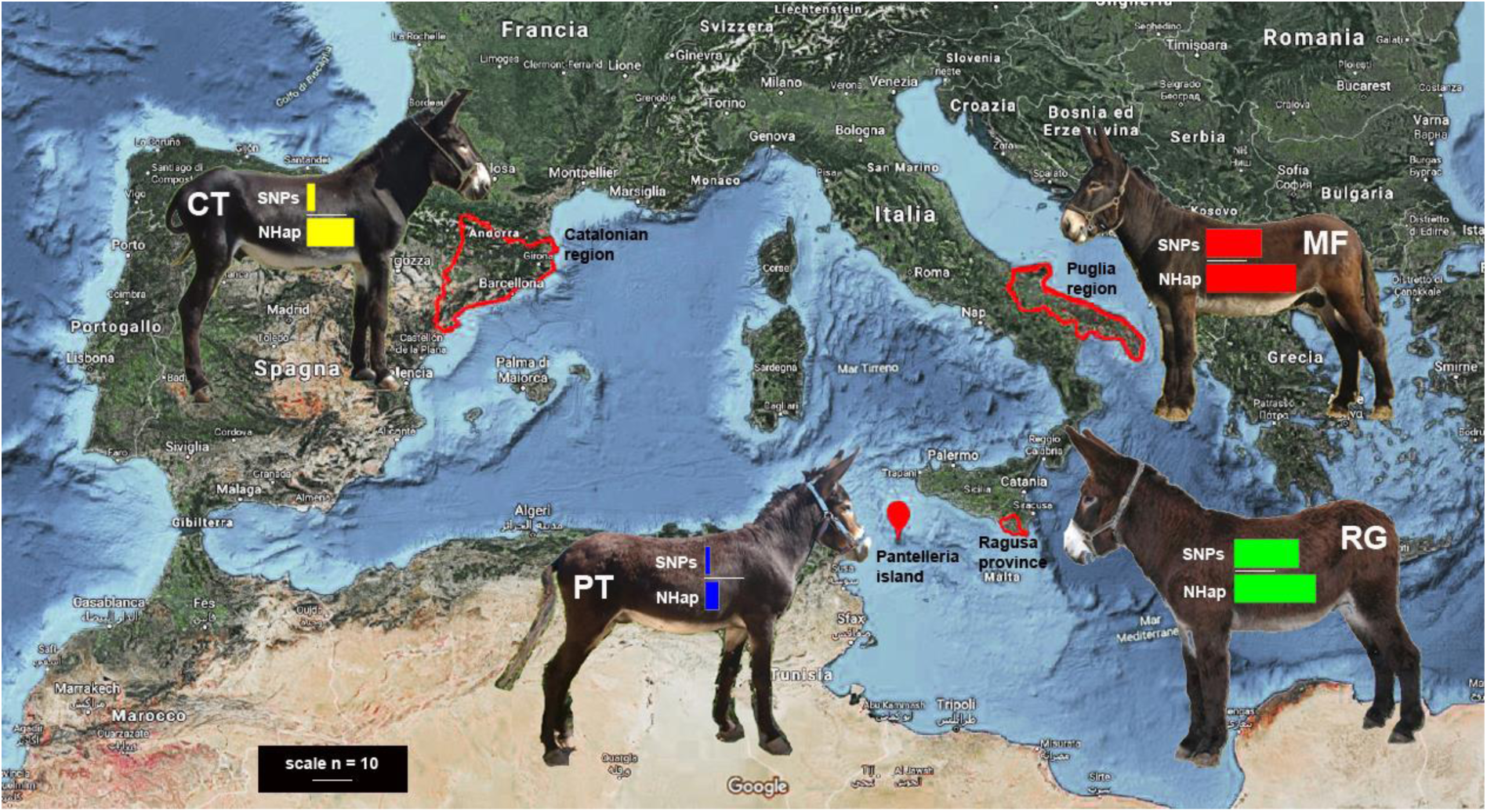
Breeds distribution and principal genetic characteristics. The red perimeter pinpoints the original Mediterranean distribution of the four donkey breed investigated: Martina Franca (MF); Ragusano (RG); Pantesco (PT); Catalonian (CT). Number of haplotypes and SNPs are indicated by horizontal histogram bar, scale bar is n = 10.

### Molecular characteristics

The highest haplotype diversity (HapD) values among all tested breeds was in MF followed by RG, the lower variability was in PT and CT (Tab. 2). The breed genetic diversity evaluated by the nucleotide diversity value (π) per breed is in Tab. 2, for the entire group of breeds is 0.128, within subpopulations is 0.098 and the mean inter-population evolutionary π is 0.03.

**Table 2.** D-loop nucleotide polymorphisms and molecular diversity indices per breed tested in the study.

| Breed | n | NHap | HapD | SNPs | $\pi$ | k ± SD | | | |
| --- | --- | --- | --- | --- | --- | --- | --- | --- | --- |
|  |  |  |  |  |  | T | C | A | G |
| Martina Franca (MF) | 27 | 22(5s) | 0.130±.007 | 13 | 0.162 | 27.1±.32 | 29.5±.39 | 33.3±.35 | 10.1±.37 |
| Ragusano (RG) | 22 | 22(3s) | 0.112±.008 | 15 | 0.132 | 27.4±.93 | 28.9±.68 | 33.1±.66 | 10.6±1.26 |
| Pantesco (PT) | 8 | 6(3s) | 0.025±.007 | 1 | 0.025 | 27.6±1.02 | 28.8±.61 | 33.2±1.11 | 10.5±1.41 |
| Catalonian (CT) | 19 | 14(8s) | 0.037±.008 | 2 | 0.038 | 27.0±.15 | 29.8±.50 | 33.5±.29 | 9.7±.15 |
| mongrel | 1 | 1 |  | 1 |  |  |  |  |  |
| ALL | 77 | 56 |  | 33 | 0.128 |  |  |  |  |

s is for shared haplotypes.
In the dataset we have a mongrel with its own haplotype.
n number of sampled individuals per breed. Nhap the number of haplotypes resulted in each breed, in parenthesis common Hap. HapD Haplotype diversity with its standard deviation; SNPs the number of polymorphic sites; $\pi$ nucleotide diversity; k average number of nucleotide differences with its standard deviation. The analyses were conducted in MEGA7.

The single nucleotide polymorphisms (SNPs) identification was performed on the absolute number of mutations found in each single sequence. According to the position towards the reference sequence, the type of mutation and the fraction in which it appears each SNPs was characterized (Fig.2).

**Fig.2.**
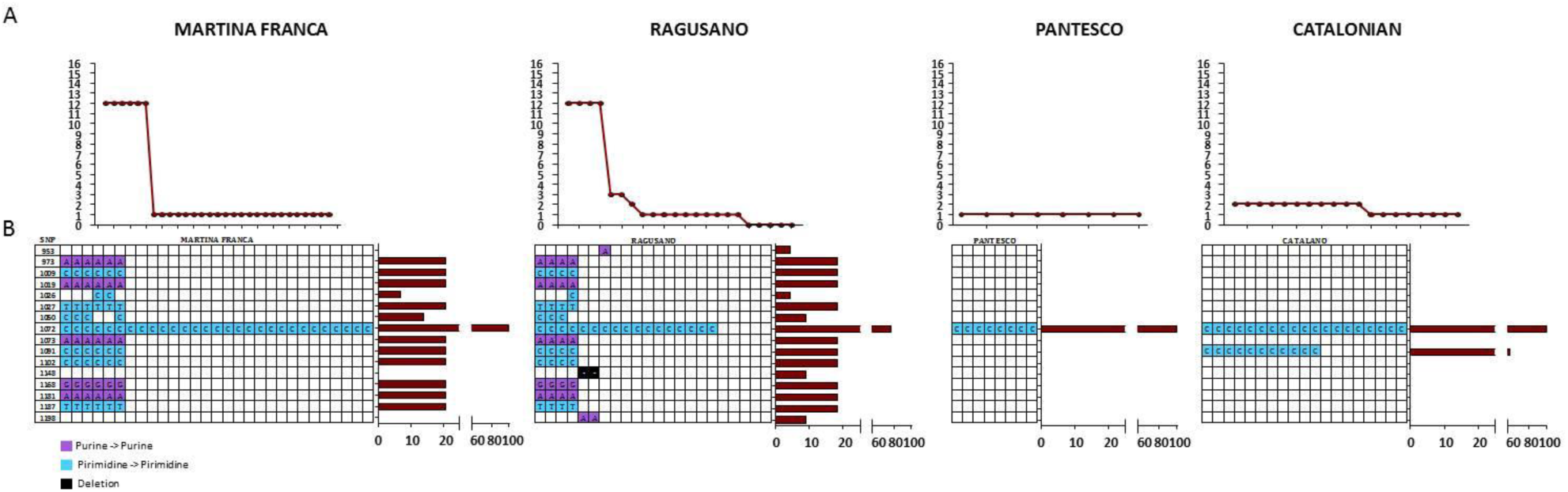
SNPs analysis performed on mitochondrial DNA isolated from listed donkey breeds. A) Absolute number of mutations found in each sample by Sanger sequencing. B) Every SNP has been characterized according to the position on the reference sequence, the kind of mutation, and the fraction of samples in which is reported (brown histograms), (MEGA7).

The nucleotide frequencies per breed is in Tab. 3, multivariate test of significance return no differences in nucleotide composition among breeds (p=.98).

**Tab. 3.** Nucleotide frequencies per breed, multivariate test of significance return no differences in nucleotide composition among breeds (p=.98).

|  | A | T(U) | C | G |
| --- | --- | --- | --- | --- |
| **MF** | 38.42 (2.46) | 31.51 (2.57) | 21.12 (1.30) | 8.95(2.72) |
| **RG** | 38.28 (3.76) | 31.58 (2.94) | 21.12 (0.87) | 9.02 (3.43) |
| **CT** | 38.72 (1.13) | 31.63 (0.95) | 21.48 (0.71) | 8.16 (0.65) |
| **PT** | 39.1 (0.61) | 31.58 (0.43) | 21.43 (0.57) | 7.89 (0.38) |
| **all** | 38.37 (2.81) | 31.62 (2.43) | 21.18 (1.02) | 8.83 (2.74) |

In parenthesis standard deviation ± SD. The analysis was conducted in MEGA7.

In Table 4 the maximum composite likelihood estimates of the nucleotide substitution pattern per breed, positions containing gaps and missing data were eliminated according to the literature (Kumar et al. 2016).

**Tab. 4.** Maximum Composite Likelihood Estimate of the Pattern of Nucleotide Substitution. Each entry shows the probability of substitution (r) from one base (row) to another base (column). Rates of different transitional (A x G e C x T) substitutions are shown in bold and those of transversional (G x T e A x C) substitutions are shown in italics.

|  | A | T | C | G |
| --- | --- | --- | --- | --- |
| MF transition/transversion rate ratios purines 4.617, pyrimidines 1.391; overall transition/transversion 1.008 |  |  |  |  |
| A | - | 6.41 | 4.29 | **8.4** |
| T | 7.81 | - | **5.97** | 1.82 |
| C | 7.81 | **8.91** | - | 1.82 |
| G | **36.06** | 6.41 | 4.29 | - |
| RG transition/transversion rate ratios purines 3.872, pyrimidines 0.95; overall transition/transversion 0.791 |  |  |  |  |
| A | - | 7.29 | 4.88 | **8.06** |
| T | 8.84 | - | **4.63** | 2.08 |
| C | 8.84 | **6.93** | - | 2.08 |
| G | **34.21** | 7.29 | 4.88 | - |
| CT transition/transversion rate ratios purines 0.985, pyrimidines 0; overall transition/transversion 0.125 |  |  |  |  |
| A | - | 12.85 | 8.73 | **3.27** |
| T | 15.73 | - | **0** | 3.32 |
| C | 15.73 | **0** | - | 3.32 |
| G | **15.49** | 12.85 | 8.73 | - |
| PT transition/transversion rate ratios purines, pyrimidines and overall transition/transversion is 0 |  |  |  |  |
| A | - | 15.79 | 10.71 | **0** |
| T | 19.55 | - | **0** | 3.95 |
| C | 19.55 | **0** | - | 3.95 |
| G | **0** | 15.79 | 10.71 | - |

For simplicity, the sum of r values is made equal to 100. All positions containing gaps and missing data were eliminated. Evolutionary analyses were conducted in MEGA7.

### Population analyses

The higher mean sequence distance within and between breed sequences are reported in Tab. 5.

**Tab. 5.** Estimates of base composition bias difference between breed sequences.

| Composition Distance | Average within groups |
| --- | --- |
| MF | 0.289 |
| RG | 0.521 |
| CT | 0.044 |
| PT | 0.020 |
| Average between groups |  |
| MF x CT | 0.162 |
| MF x RG | 0.387 |
| MF x PT | 0.158 |
| RG x CT | 0.276 |
| RG x PT | 0.278 |
| CT x PT | 0.031 |

The analysis were conducted in MEGA7.

The evolutionary relationships and relative abundance of haplotypes per breed is calculated by using the UPGMA and Maximum Composite Likelihood method (Fig. 3). The most represented haplotypes are seven: 51 is common in all breeds; 22, 36 and 37 found in MF are shared with PT and/or RG; 30 distinctive of MF is also found in CT; 40 and 53 are characteristics of CT.

**Fig.3.**
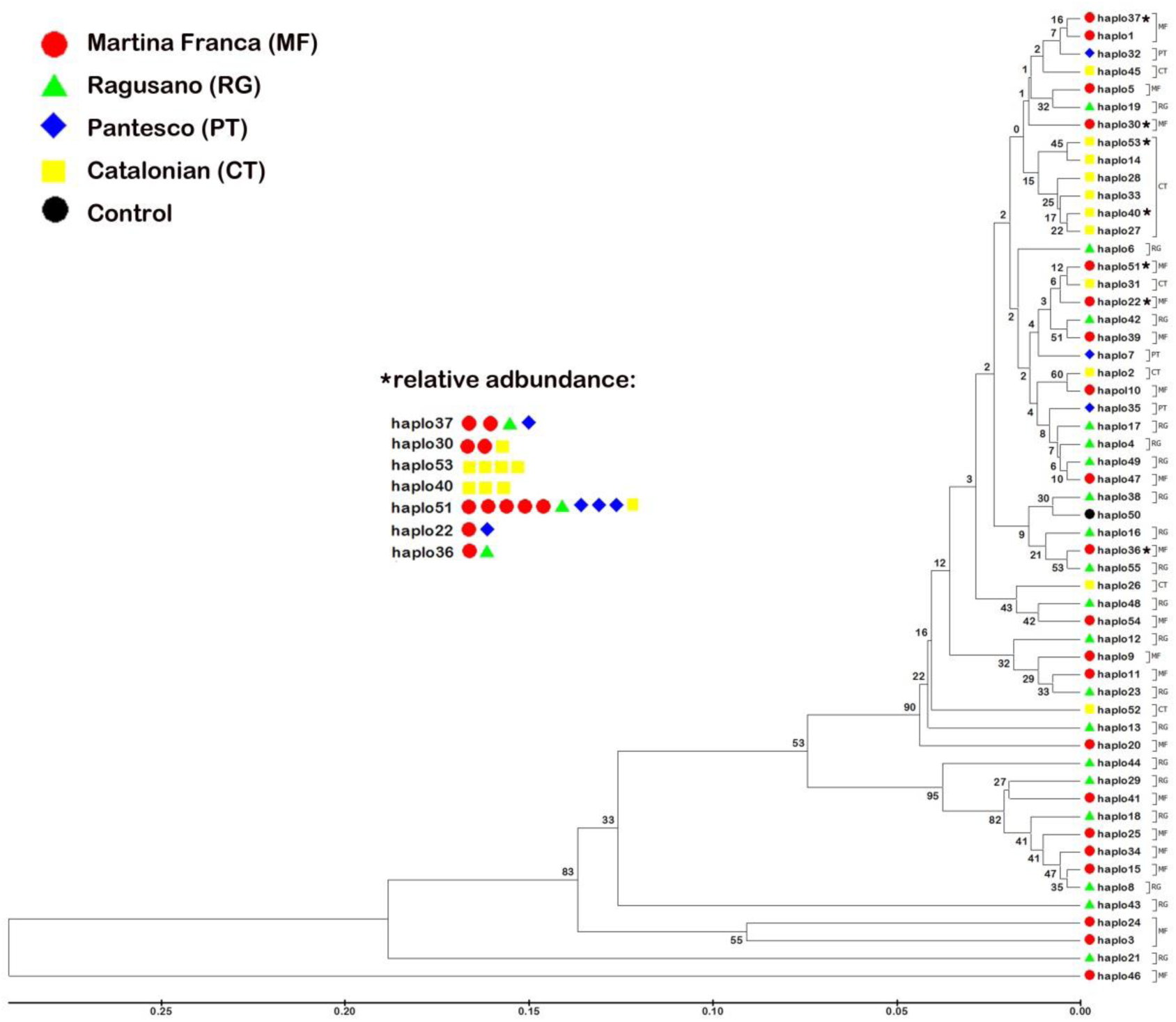
Evolutionary relationships of haplotypes. The evolutionary history was inferred using the UPGMA method. The optimal tree with the sum of branch length = 1.85287086 is shown. The percentage of replicate trees in which the associated taxa clustered together in the bootstrap test (500 replicates) are shown next to the branches. The tree is drawn to scale, with branch lengths in the same units as those of the evolutionary distances used to infer the phylogenetic tree. The evolutionary distances were computed using the Maximum Composite Likelihood method and are in the units of the number of base substitutions per site. Evolutionary analyses were conducted in MEGA7.

### Origin and phyletic relationship

The phyletic relationships among the 55 haplotypes identified is calculated through the Median-joining Network. The haplotypes of each breed are identified by the color code, the abundance by the relative size of the symbol and the diffusion among the breeds with the pie division of the various colors (Fig.4).

**Fig.4.**
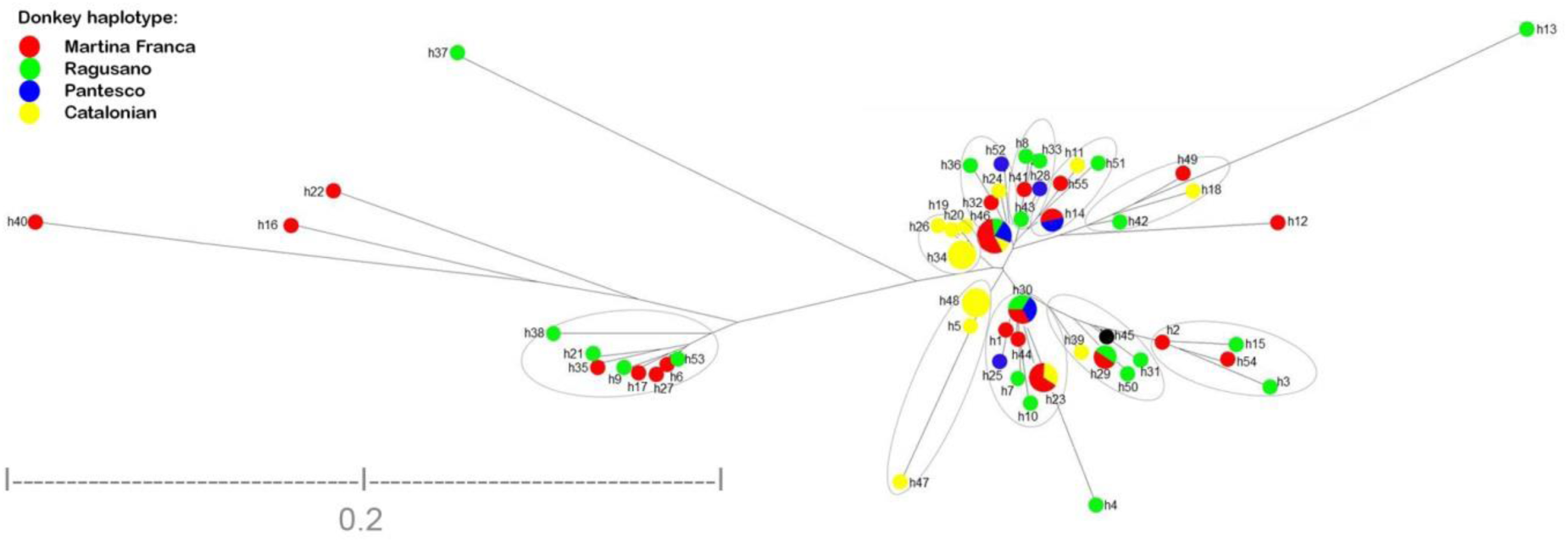
The median-joining network. It is based on 350 bp of the mitochondrial D-Loop representing 77 donkeys within 55 haplotypes (DARwin 6.0). Each breed was shown by color and the proportions of different breed for each haplotype were shown. Reference mongrel samples is represented by black sphere.

**Fig.5.**
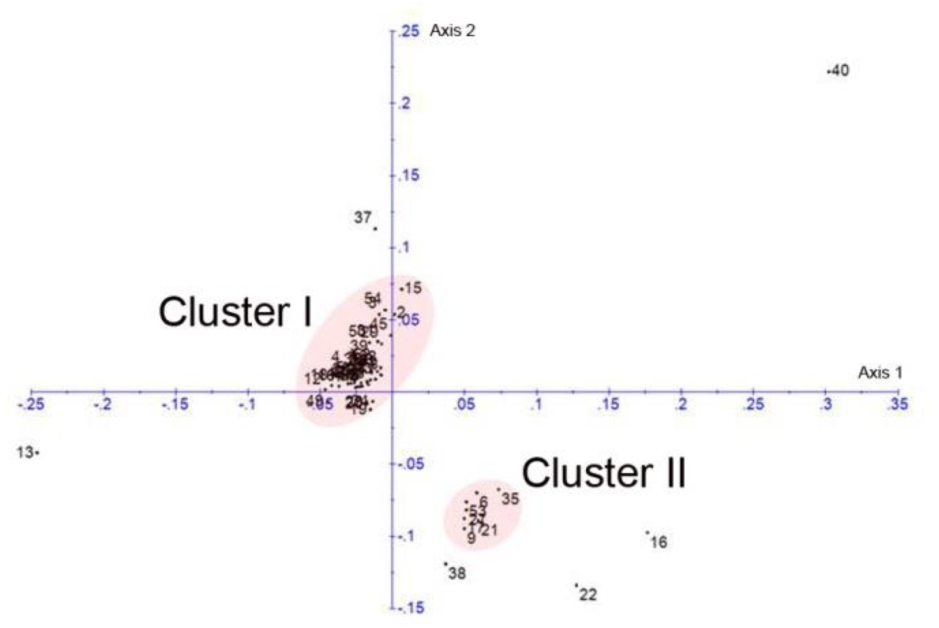
Principal Coordinates Analysis (PCoA). The PCoA plot of the two first axes based upon the dissimilarity matrix according to Kimura (1980), (DARwin 6.0). We found two different clusters, cluster I and cluster II includes most haplogroups, external haplotypes are: MF 22, 37 and 40; RG 13, 16 and 38.

The Principal Coordinates (PCoA) analysis, based on the dissimilarity matrix, return two different clusters, I and II, grouping most haplogroups. However, several haplotypes are external to both clade and are MF 22, 37 and 40; RG 13, 16 and 38.

## Discussion

### Breed analysis

Donkey breed represents a fascinating model of domesticated biodiversity; thus a number of studies address donkey pedigree or genetic analysis. Pedigree study and reconstruction, usually, deficit in genetic correspondence (Gutiérrez et al, 2005; Cecchi et al, 2006; Rizzi et al, 2011, Navas et al, 2017), while genetic ones frequently lack lineage information (Bordonaro et al, 2012; Cinar Kul et al, 2016; Cozzi et al, 2017).

Overall, studies on pedigree in Italian (MF, Amiata) and Spanish (CT, Andalusian, Miranda) donkey found a dramatic loss of genetic variation, essentially due to the high rate of inbreeding (Gutiérrez et al, 2005; Cecchi et al, 2006; Rizzi et al, 2011; Navas et al, 2017). However, the weakness of pedigree completeness, the occurrence of a bottleneck event (e.g. MF, in 1980), may cause over or underestimation of the data and consequentially could affect the breeding strategy (Rizzi et al, 2011; Navas et al, 2017). To overcome this bias, in the present study genetic analysis were performed on subject with Certificate of Origin, produced by official authorized breeding center. This approach allowed us to link mtDNA data to pedigree records per population and breed, in order to: identify the same maternal descendant, select individual with the presumed higher genetic variability and preserve biodiversity. A further bias in studies on breed: population, genetic structure, variability, differences, robustness, average relatedness, inbreeding, co-ancestry, degree of non-random mating and origin (e.g. Aranguren-Mendez et al, 2004; Ivanković et al 2009; Cinar Kul et al, 2016), is from unbalanced sample size comparisons, e.g. of fivefold of difference (Cozzi et al, 2017; Bordonaro et al, 2012). Conversely, here an equal number of samples per breed were collected, consequently all analysis refers to a balanced sample. Higher genetic variability in MF and RG were found, which disagrees with work-based on pedigree (Rizzi et al, 2011). RG and MF are widely used in the farm and still have a natural random mating that is, we think, the source of variability. The lower variability is in PT, as expected, because of its isolation. In the PT Certificate of Origin eight distinct maternal lineages are attested, which are in agreement with the statement that the breed recovery starting from a small nucleus of nine founders (Bordonaro et al, 2012).

Furthermore, by looking at the number of haplotypes the PT genetic robustness is dramatically lower than predictable by the pedigree information lonely. On the contrary, low variability found in CT is unpredicted, because of the number of individuals in population commonly distributed in the largest region of Catalonia. This phenomenon, possibly, could be related to breeding program rather than the number of subjects and dimension of the homeland. Consequently, a potential bottleneck is produced by human artificial selection, which caused lost in variability according to previous studies (Gutiérrez et al, 2005; Navas et al, 2017).

Therefore, in the light of variability, robustness and the degree of non-random mating decrease with average relatedness, inbreeding and co-ancestry increase in breed genetic structure, thus a new program of reproduction with multiple approaches is needed.

### Molecular characteristics

The molecular characteristics analyzed shows distinctive nucleotide frequency among breeds, which are in line with literature (Xu et al, 1996; Luo et al 2011). The transition/transversion rate ratios is greater for purines, transition/transversion per breed is decreasing from MF to RG and CT, which is discordant to other study on Italian donkey breed (Cozzi et al, 2017). The nucleotide diversity (π) values is in line with the study on CT (Aranguren-Mendez et al, 2004) while not with the analysis reported on MF and RG by Cozzi et al (2017).

Furthermore, the highest haplotypes diversity HapD were found in MF followed by RG, while the lower variability was in CT followed by PT. The differences with previous study on Italian breed could be related to the following biases in Cozzi et al (2017): *i.* lack information about genealogy with consequent uncertain breed origin; *ii.* unbalanced sampling between breeds with a high contribution of Asinara and Sardo donkeys 74%, both from Sardinia island; further, *iii.* the analysis do not distinguish among breeds.

Overall, molecular indices show greater genetic variability among the Italian breeds than Spanish, according to conclusion of previous research (Aranguren-Mendez et al, 2004; Cozzi et al, 2017; Navas et al, 2017).

### Population analyses

The population analysis showed higher difference inside RG, followed by MF, lasting CT and PT. RG population genetic structure is analogous to the maternal landscape of the highly heterogeneous large Balkan donkey population, with a genetic structure more complex than previously thought (Stanisic et al, 2017). The Serbian donkey population is highly genetically diverse despite the severe population decline, probably due to the introgression of other related breeds, which support the population heterogeneity (Stanisic et al, 2017). A similar action was probably carried out for RG while not in MF. CT and PT are consistent with previous paragraph and with the outcome of a study on microsatellites genetic variability confirming lower variability in PT comparing RG and another Sicilian breed (the ‘Grigio Siciliano’ GS; Bordonaro et al, 2012). Accordingly, the genetic variability observed in PT, RG and GS (Bordonaro et al, 2012) was lower than that reported in five Spanish breeds (Aranguren-Mendez et al, 2001) and three Croatian breeds (Ivanković et al, 2002), but higher than that observed in the Amiata donkey from Italy (Ciampolini et al, 2007) and in Chinese breeds (Chen et al, 2006).

The comparison between breeds shows the higher difference between MF and RG, while lower with PT and CT comparing, a similar difference are between RG with PT and CT, finally between the PT and CT are almost similar. These results have previously experienced in two different studies on Balkan donkey with different interpretations. In the first one, a lack of correspondence between geographical areas and maternal genetic structure was found, such as the differentiation between the Balkan donkey and the African Burkina Faso donkeys outgroup also was low, the authors report a difficulties to trace routes of expansion in donkey, consequently they suggest the hypothesis of a very quick spread of the species after domestication (Pérez-Pardal et al, 2014). In the second paper, a detailed study was assessed on three Balkan breeds: Istrian (IS), north Adriatic (NA) and Littoral-Dinaric (LD) donkey populations, suggesting IS a unique breed that interfered in LD by sporadic migration events, further NA and LD have similar genetics (Ivankovic et al, 2002). By mediating this results to our paper similar effects of migration were assessed by MF in CT and PT, accordingly with historical reconstruction.

### Origin and phyletic relationship

The well-established identification of two main lineages and the probable existence of another unrecognized extinct wild ancestor in domestic donkeys is believed to be the result of separate domestication events (Uerpmann, 1991; Vila, 2002; Rossel et al, 2008; Beja-Pereira et al, 2004; Chen et al, 2006; Kimura et al, 2011; Kefena et al, 2014). Therefore, in donkey, such as in dog (Verginelli et al, 2005), genetic data support the hypothesis of the multi-centric origin of breed. According to this emerging theory is the identification of a potential new clade, unique of MF and RG. In recent study a similar hypothesis has been suggested (Cozzi et al, 2017). Accordingly, in Croatia and in Serbia donkey three haplotype groups were found (Ivankovic et al, 2002) with distinct nuclear gene pool (Stanisic et al 2017). The heterogeneous genetic structure of Balkan donkey was hypothesized due to no geographical structure and consequently was difficult to trace routes of expansion in donkey (Pérez-Pardal, 2014). However, another hypothesis on the complexity of genetic relationships among Italian donkey breed and those belonging to breeds living in Mediterranean and Balkan areas (Cozzi et al, 2017) could be due to the ancestry or from the genetic makeup of the modern donkey populations. The lasting speculating hypothesis arise from our analysis suggesting a multicentric domestication phenomena coupled with multiple waves colonization and contra-colonization such occur in Spain with MF bring by Romanian empire and later in Italy by Spanish domination, which is according with a similar phenomenon suggested by Stanisic and colleague (2017).

The rising interest in management of genetic diversity in animal populations is to safeguard the widest possible genetic resources through conservation programs (Bjørnstad & Ried, 2002; Toro et al, 2003).

In the conservation of domestic breeds, the preservation of genetic capitals is crucial because they are zoo-technical form, which already have had a reduction in the original natural variability. The biodiversity of the domestic form is the result of a genetic pool derived from the interaction with semi-artificial environments, regulated by human needs and its migratory movements. As a result, genetic and phenotypic changes with respect to the wild species of origin have been slowly addressed by human needs.

The donkey conservation represents a biological problem connected to the analogy in phenotype. The hypothesis on likeness is related to genetic exchange among breeds; and/or identical origin, from Africa or Asia population; and/or equivalent climate condition; and/or same type of work.

From the molecular analyses carried out a dramatic variability lost occur in all breeds, in particular in PT and CT. Conversely, MF and RG shows higher number of haplotypes and SNPs.

Further, it emerges that the Martina Franca breed is probably the progenitor of the Pantesco and Catalano. The evaluation of the evolutionary relationships between Martinese and Ragusano is more complex similar to Balkan donkey.

In conclusion, the study carried out determined: i. the identification of D-loop mtDNA characteristic for the Martina Franca donkey and three phenotypically similar breeds; ii. the identification of different matrilines in the Martinese and other breeds; iii. the identification of biodiversity for each breed; iv. the phyletic relationships between the breeds.

Finally, this extensive study on biodiversity and phyletic relationships in Martina Franca and her like Ragusano, Pantesco and Catalano is useful for future studies on the domestication of the species.

In practice, the Martinese donkey is an important reserve of biodiversity that must necessarily be preserved in the widest possible range of its genetic heritage, avoiding, through conservation programs, errors of consanguinity for aesthetic purposes, but considering each individual animal precious for the conservation of the breed because aesthetics does not often coincide with the genetic heritage within the subject, so it is concluded that future conservation programs will necessary include in the Certificate of Origin also the genetic analysis of at least the matrilines.

## Bibliografia

1. Aranguren-Mendez J, Beja-Pereira A, Avellanet R, Dzama K, Jordana J (2004) Mitochondrial DNA variation and genetic relationships in Spanish donkey breeds (Equus asinus). J Anim Breed Genet 121:319–330.

2. Aranguren-Mendez J, Jordana J and Gomez M (2001) Genetic diversity in Spanish donkey breeds using microsatellite DNA markers. Genetics Selection Evolution 33(4):433–442.

3. Achilli A, Olivieri A, Soares P, Lancioni H, Kashani BH, Perego UA, Nergadze SG, Carossa V, Santagostino M, Capomaccio S, et al. (2012) Mitochondrial genomes from modern horses reveal the major haplogroups that underwent domestication. Proc Natl Acad Sci USA, 109(7):2449–2454.

4. Beja-Pereira, A. et al. 2004 African origins of the domestic donkey. Science 304, 1781. (doi:10.1126/science.1096008).

5. Bertolini F, Scimone C, Geraci C, Schiavo G, Utzeri VJ, Chiofalo V, Fontanesi L (2015) Next Generation semiconductor based sequencing of the donkey (*Equus asinus*) genome provided comparative sequence data against the horse genome and a few millions of single nucleotide polymorphisms. PLoS One. 10:e0131925.

6. BjØrnstad G, Ried KH (2002) Evaluation of factors affecting individual assignment precision using microsatellite data from horse breeds and simulated breed crosses. Animal Genet 33: 264–270.

7. Bordonaro S, Guastella AM, Criscione A, Zuccaro A, Marletta D. 2012. Genetic diversity and variability in endangered Pantesco and two other Sicilian donkey breeds assessed by microsatellite markers. Sci World J. 648427, 1-6 doi:10.1100/2012/648427

8. Bowling AT, Del Valle A, Bowling M: A pedigree-based study of mitochondrial D-loop DNA sequence variation among Arabian horses. Anim Genet 2000, 31(1):1–7.

9. Brown WM, George M Jr, Wilson AC (1979) Rapid evolution of animal mitochondrial DNA. Proc Natl Acad Sci U S A 76: 1967–1971.

10. Cecchi F, Ciampolini R, Ciani E, Matteoli B, Mazzanti E, Tancredi M, Presciuttini S (2006) Demographic genetics of the endangered Amiata donkey breed. Ital J Anim Sci 5:387–391.

11. Chen, S. Y., Zhou, F., Xiao, H., Sha, T., Wu, S. F. & Zhang, Y. P. 2006 Mitochondrial DNA diversity and population structure of four Chinese donkey breeds. Anim. Genet. 37, 422–431.

12. Cinar Kul B, Bilgen N, Akyuz B, Ertugrul O (2016) Molecular phylogeny of Anatolian and Cypriot donkey populations based on mitochondrial DNA and Y-chromosomal STRs. Ankara Üniv Vet Fak Derg, 63:143–149.

13. Ciampolini R, Cecchi F, Mazzanti E, Ciani E, Tancredi M, De Sanctis B (2007) The genetic variability analysis of the Amiata donkey breed by molecular data. Italian J Anim Sci. 6(1):78–80.

14. Cieslak M, Pruvost M, Benecke N, Hofreiter M, Morales A, Reissmann M, Ludwig A: Origin and History of Mitochondrial DNA Lineages in Domestic Horses. Plos One 2010, 5:12.

15. Clutton-Brock, J. 1992 Horse power: a history of the horse and the donkey in human societies. London, UK: Harvard University Press and the Natural History Museum.

16. Colli L, Perrotta G, Negrini R, Bomba L, Bigi D, Zambonelli P, Verini Supplizi A, Liotta L, Ajmone-Marsan P. 2013. Detecting population structure and recent demographic history in endangered livestock breeds: the case of the Italian autochthonous donkeys. Anim Genet. 44:69–78.

17. Cothran EG, Juras R, Macijauskiene V: Mitochondrial DNA D-loop sequence variation among 5 maternal lines of the Zemaitukai horse breed. Genet Mol Biol 2005, 28(4):677–681.

18. Cozzi MC, Valiati P, Cherchi R, Gorla E, Prinsen RTMM, Longeri M, Bagnato A and Strillacci MG (2017) Mitochondrial DNA genetic diversity in six Italian donkey breeds (Equus asinus). Mitochondrial DNA part A 1-10.

19. George MJ, Ryder OA (1986) Mitochondrial DNA evolution in the genus Equus. Mol Biol Evol 3(6):535–546

20. Groves C, Grubb P (2011) Ungulate taxonomy: Johns Hopkins University Press.

21. Guastella AM, Zuccaro A, Criscione A, Marletta D, Bordonaro S (2011) Genetic Analysis of Sicilian Autochthonous Horse Breeds Using Nuclear and Mitochondrial DNA Markers. J Heredity 102(6):753–758.

22. Guastella AM, Zuccaro A, Bordonaro S, Criscione A, Marletta D, D’Urso G (2007) Genetic diversity and relationship among the three autochthonous Sicilian donkey populations assessed by microsatellite markers. Italian J Anim Sci 6(1):143.

23. Gutiérrez JP, Marmi J, Goyache F, Jordana J (2005) Pedigree information reveals moderate to high levels of inbreeding and a weak population structure in the endangered Catalonian donkey breed. J Anim Breed Genet 122:378–386.

24. Hutchison CA 3rd, Newbold JE, Potter SS, Edgell MH: Maternal inheritance of mammalian mitochondrial DNA. Nature 1974, 251(5475):536–538.

25. Ishida N, Hasegawa T, Takeda K, Sakagami M, Onishi A, et al. (1994) Polymorphic sequence in the D-loop region of equine mitochondrial DNA. Anim Genet 25: 215–221.

26. Ivanković A, Ramljak J, Konjačić M, Kelava N, Dovč P, Mijić P (2009) Mitochondrial D-loop sequence variation among autochthonous horse breeds in Croatia. Czech J. Anim. Sci. 54(3):101–111.

27. Ivankovic A, Kavar T, Caput P, Mioc B, Pavic V, Dovc P (2002) Genetic diversity of three donkey populations in the Croatian coastal region. Animal Genetics 33(3):169–177.

28. Jansen T, Forster P, Levine MA, Oelke H, Hurles M, Renfrew C, Weber J, Olek K: Mitochondrial DNA and the origins of the domestic horse. Proc Natl Acad Sci USA 2002, 99(16):10905–10910.

29. Kavar T, Dovc P (2008) Domestication of the horse: Genetic relationships between domestic and wild horses. Livest Sci 116(1–3):1–14.

30. Kefena E, Dessie T, Tegegne A, Beja-Pereira A, Yusuf Kurtu M, Rosenbom S, Han JL. (2014) Genetic diversity and matrilineal genetic signature of native Ethiopian donkeys (*Equus asinus*) inferred from mitochondrial DNA sequence polymorphism. Livestock Sci 167:73–79.

31. Kimura B, Marshall FB, Chen S, Rosenbom S, Moehlman PD, Tuross N, Sabin RC, Peters J, Barich B, Yohannes H, Kebede F, Teclai R, Beja-Pereira A, Mulligan CJ (2011). Ancient DNA from Nubian and Somali wild ass provides insights into donkey ancestry and domestication. Proceedings of the Royal Society B: Biological Sciences, 278:50–7.

32. Kivisild T, Reidla M, Metspalu E, Rosa A, Brehm A, Pennarun E, Parik J, Geberhiwot T, Usanga E, Villems R: Ethiopian mitochondrial DNA heritage: tracking gene flow across and around the gate of tears. Am J Hum Genet 2004, 75(5):752–770.

33. Kruger K, Gaillard C, Stranzinger G, Rieder S (2005) Phylogenetic analysis and species allocation of individual equids using microsatellite data. Journal of Animal Breeding and Genetics 122: 78–86.

34. Kumar S, Stecher G, Tamura K (2016). MEGA7: Molecular Evolutionary Genetics Analysis Version 7.0 for Bigger Datasets. Mol. Biol. Evol. 33(7):1870–1874.

35. Kumar S., Stecher G., and Tamura K. (2015). MEGA7: Molecular Evolutionary Genetics Analysis version 7.0 for bigger datasets. Molecular Biology and Evolution.

36. Lindsay EH, Opdyke ND, Johnson NM (1980) Pliocene dispersal of the horse Equus and late Cenozoic mammalian dispersal events. Nature 287:135–138

37. Lippold S, Matzke NJ, Reissmann M, Hofreiter M: Whole mitochondrial genome sequencing of domestic horses reveals incorporation of extensive wild horse diversity during domestication. Bmc Evol Biol 2011, 11:328.

38. Luo Y, Chen Y, Liu FY, Jiang CH, Gao YQ (2011) Mitochondrial genome sequence of the Tibetan wild ass (Equus kiang). Mitochondrial DNA 22: 6–8.

39. MacFadden BJ (2005) Fossil horses - Evidence for evolution. Science 307: 1728–1730.

40. Matassino D, Cecchi F, Ciani F, Incoronato C, Occidente M, Santoro L, Ciampolini R. 2014. Genetic diversity and variability in two Italian autochthonous donkey genetic types assessed by microsatellite markers. Italian. J Anim Sci. 13:3028.

41. Matisoo-Smith E, Robins J: Mitochondrial DNA evidence for the spread of Pacific rats through Oceania. Biol Invasions 2009, 11(7):1521–1527.

42. Mazzatenta A, Carluccio A, Robbe D, Di Giulio C, Cellerino A (2017). The companion dog as a unique translational model for aging. Semin Cell Dev Biol.; 70:141–153.

43. Mirol PM, Garcia PP, Vega-Pla JL, Dulout FN: Phylogenetic relationships of Argentinean Creole horses and other South American and Spanish breeds inferred from mitochondrial DNA sequences. Anim Genet 2002, 33(5):356–363.

44. Navas FJ, Jordana J, León JM, Barba C, Delgado JV (2017) A model to infer the demographic structure evolution of endangered donkey populations. Animal, 1–10.

45. Pérez-Pardal L, Grizelj J, Traoré A, Cubric-Curik V, Arsenos G, Dovenski T, Marković B, Fernández I, Cuervo M, Alvarez I, Beja-Pereira A, Curik I, Goyache F (2014) Lack of mitochondrial DNA structure in Balkan donkey is consistent with a quick spread of the species after domestication. Anim Genet 45(1):144–7.

46. Piganiol AFG La conquête romaine 1927 Edité par Félix Alcan, Paris.

47. Rand DM (2001) The units of selection on mitochondrial DNA. Annu Rev Ecol Syst 32: 415–448.

48. Rizzi R, Tullo E, Cito AM, Caroli A and Pieragostini E (2011) Monitoring of genetic diversity in the endangered Martina Franca donkey population. J Anim Sci, 89:1304–1311

49. Rossel, S., Marshall, F., Peters, J., Pilgram, T., Adams, M. D. & O’Connor, D. 2008 Domestication of the donkey: timing, processes and indicators. Proc. Natl Acad. Sci. USA 105, 3715–3720. (doi:10.1073/pnas.0709692105)

50. Simpson GG (1951) Horses: the story of the horse family in the modern world and through sixty million years of history. Oxford University Press, New York.

51. Stanisic LJ, Aleksic JM, Dimitrijevic V, Simeunovic P, Glavinic U, Stevanovic J, Stanimirovic Z (2017) New insights into the origin and the genetic status of the Balkan donkey from Serbia. Anim Genet 48(5):580–590.

52. Stoneking M, Sherry ST, Redd AJ, Vigilant L. (1992). New approaches to dating suggest a recent age for the human mtDNA ancestor. Philos Trans R Soc Lond B Biol Sci. 29;337(1280):167–75.

53. Tamura K., Battistuzzi FU, Billing-Ross P, Murillo O, Filipski A, and Kumar S. (2012). Estimating Divergence Times in Large Molecular Phylogenies. Proceedings of the National Academy of Sciences 109:19333–19338.

54. Tamura K., Nei M., and Kumar S. (2004). Prospects for inferring very large phylogenies by using the neighbor-joining method. Proceedings of the National Academy of Sciences (USA) 101:11030–11035.

55. Toro MA, Barragan C, Ovilo C (2003) Estimation of genetic variability of the founder population in a conservation scheme using microsatellites. Ani Genet 34: 226–228.

56. Uerpmann, H.-P. 1991 Equus africanus in Arabia. In Equids in the Ancient World (eds R. H. Meadow & H.-P. Uerpmann), pp. 12–33. Wiesbaden: Ludwig Reichert Verlag

57. Verginelli F, Capelli C, Coia V, Musiani M, Falchetti M, Ottini L, Palmirotta R, Tagliacozzo A, De Grossi Mazzorin I, Mariani-Costantini R. (2005). Mitochondrial DNA from prehistoric canids highlights relationships between dogs and South-East European wolves. Mol Biol Evol.; 22(12):2541–51.

58. Vigilant L, Pennington R, Harpending H, Kocher TD, Wilson AC (1989). Mitochondrial DNA sequences in single hairs from a southern African population. Proc Natl Acad Sci U S A; 86(23):9350–4.

59. Vilá C, Leonard JA, Beja-Pereira A: Genetic Documentation of Horse and Donkey Domestication. In Documenting Domestication. edited by Zeder MA, Bradley DG, Emshwiller E, Smith BD.; 2006:342–353.

60. Vila C, Leonard JA, Gotherstrom A, Marklund S, Sandberg K, Liden K, Wayne RK, Ellegren H: Widespread origins of domestic horse lineages. Science 2001, 291(5503):474–477.

61. Vila, E. 2002 Data on equids from late fourth and third millennium sites in Northern Syria. In Equids in time and space (ed. M. Mashkour), pp. 101–123. Oxford, UK: Oxbow Books.

62. Wallace DC (2007) Why do we still have a maternally inherited mitochondrial DNA? Insights from evolutionary medicine. Annu Rev Biochem 76: 781–821.

63. Xu XF, Gullberg A, Arnason U (1996) The complete mitochondrial DNA (mtDNA) of the donkey and mtDNA comparisons among four closely related mammalian species-pairs. Journal of Molecular Evolution 43: 438–446.

64. FAO report http://dad.fao.org/

